# Optimizing therapeutic outcomes with Mechanotherapy and Ultrasound Sonopermeation in solid tumors

**DOI:** 10.1101/2024.11.28.625828

**Authors:** Marina Koutsi, Triantafyllos Stylianopoulos, Fotios Mpekris

**Affiliations:** Cancer Biophysics Laboratory, Department of Mechanical and Manufacturing Engineering, University of Cyprus, Nicosia, Cyprus

**Keywords:** mathematical model, drug delivery, tumor microenvironment, nano-immunotherapy

## Abstract

Mechanical solid stress plays a pivotal role in tumor progression and therapeutic response. Elevated solid stress compresses intratumoral blood vessels, leading to hypoperfusion, and hypoxia, which impair oxygen and drug delivery. These conditions hinder the efficacy of drugs and promote tumor progression and treatment resistance compromising therapeutic outcomes. To enhance treatment efficacy, mechanotherapeutics and ultrasound sonopermeation have been developed to improve tumor perfusion and drug delivery. Mechanotherapy aims to reduce tumor stiffness and mechanical stress within tumors to normal levels leading to decompression of vessels while simultaneously improving perfusion. On the other hand, ultrasound sonopermeation strategy focuses on increasing non-invasively and transiently tumor vessel wall permeability to boost perfusion and thus, improve drug delivery. Within this framework and aiming to replicate published experimental data in silico, we developed a mathematical model designed to derive optimal conditions for the combined use of mechanotherapeutics and sonopermeation, with the goal of optimizing efficacy of nano-immunotherapy. The model incorporates complex interactions among diverse components that are crucial in the multifaceted process of tumor progression. These components encompass a variety of cell populations in tumor, such as tumor cells and immune cells, as well as components of the tumor vasculature including endothelial cells, angiopoietins, and the vascular endothelial growth factor. A comprehensive validation of the predictions generated by the mathematical model was carried out in conjunction with published experimental data, wherein a strong correlation was observed between the model predictions and the actual experimental measurements of critical parameters, which are essential to reinforce the overall accuracy of the mathematical framework employed. In addition, a parametric analysis was performed with primary objective to investigate the impact of various critical parameters that influence sonopermeation. The analysis provided optimal guidelines for the use of sonopermeation in conjunction with mechanotherapy, that contribute to identify optimal conditions for sonopermeation.

## 1 Introduction

Solid tumors are complex biological entities that are composed not solely of malignant cellular populations but also include various additional components, such as stromal cells and extracellular matrix (ECM), which collectively form the tumor microenvironment (TME) [1–5]. In numerous instances of tumor pathophysiology, particularly in highly desmoplastic cancers such as various sarcoma subtypes, the TME becomes fibrotic as the tumor proliferates. This fibrosis is indicative of an augmented synthesis of ECM elements, predominantly collagen and hyaluronan, which contribute to the development of a tumor mass with elevated stiffness, thereby impacting its mechanical properties [6]. The high density of cancer cells, stromal components, and extracellular matrix elements, coupled with the accelerated growth of the tumor at the expense of surrounding host tissue, generates mechanical forces referred to as solid stress, which manifest both within the tumor and between the tumor and surrounding host tissue [7, 8]. The role of mechanical solid stresses is critical in influencing the progression of tumors as well as the efficacy of therapeutic interventions. The elevation of solid stress can induce compression of intratumoral blood vessels and ultimately result in their collapse and dysfunction, thereby compromising the delivery of oxygen and nutrients to the tumor mass, leading to hypoperfusion and hypoxia [9–11]. Hypoperfusion can severely hinder the effective intratumoral delivery of therapeutics administered systemically, while the prevailing hypoxic environment can significantly enhance tumor progression and confer resistance to treatment through a variety of mechanisms [12].

A therapeutic strategy aimed at decompressing vessels while simultaneously enhancing the perfusion levels within tumor tissues involves the application of mechanotherapeutics, which are intended to reduce tumor stiffness and the mechanical forces within tumors to normal levels [13]. This is achieved by specifically targeting ECM components, such as collagen and hyaluronan, or Cancer-Associated Fibroblasts (CAFs), thereby reopening compressed blood vessels and improving both the perfusion and distribution of therapeutic drugs within the TME [14–22]. It was demonstrated both mathematically and experimentally that the mechanotherapeutic agent tranilast, typically utilized as an anti-fibrotic drug, employs a stress alleviation strategy that enhances the functional vascular density by decompressing blood vessels and improving the delivery of therapeutic agents [16, 23]. Ketotifen, an antihistamine medication, it has been effectively demonstrated that possesses a dual functionality, whereby it serves as both a mechanomodulator and an immunomodulator within the TME specifically in sarcomas [6, 24, 25]. Nevertheless, it is important to note that the efficacy of mechanotherapeutics is somewhat limited, as they are only able to decompress a fraction of the compressed blood vessels within the tumor, rather than achieving comprehensive decompression of all vessels [19].

Ultrasound sonopermeation is a method to enhance drug delivery in solid tumors that utilizes ultrasound in combination with microbubbles. Sonopermeation can enhance transiently the permeability of vessel walls, thereby improving the delivery of therapeutic agents. This method has shown promise in overcoming biological barriers and improve drug delivery to tumors [26, 27]. It has been also found that sonopermeation can reduce intratumoral solid stress and thus improve perfusion; however, the fundamental mechanisms governing this phenomenon remain inadequately elucidated [28, 29]. Indeed, the use of ultrasound in the presence of microbubbles has exhibited enhanced therapeutic efficacy when compared to traditional nano- and chemo-therapeutic agents [27, 30–34]. Interestingly, we provided evidence that mechanotherapy can be combined with sonopermeation and that the two strategies can have synergistic effects on improving therapeutic efficacy [6].

Up to now, there has been limited research on mathematical modeling related to sonopermeation and drug delivery for the treatment of solid tumors. Few mathematical models have been formulated to investigate the impact of microbubbles within blood vessels [35, 36], often overlooking perfusion and drug delivery challenges posed by the TME and lacking experimental validation. Recently, a more detailed mathematical model has been developed that elucidates the mechanisms by which low-intensity ultrasound serves to inhibit the proliferation and expansion of cancer stem cells [37, 38]. In the present study, we developed a mathematical model for the study of the combined effects of mechanotherapy and sonopermeation, building on our previous studies [23, 39–42]. More precisely, the newly developed model has been designed to derive optimal conditions for the combined use of mechanotherapeutics and sonopermeation, by integrating the crucial effects of sonopermeation, alongside the impact of the mechanotherapeutic ketotifen on the tumor microenvironment. Furthermore, a comparison of model predictions with experimental data was carried out to substantiate the predictions generated by the mathematical model. Finally, an extensive parametric analysis was performed with the primary objective of investigating the impact of various critical parameters that significantly influence the effects of sonopermeation, which in turn aims to enhance our understanding on the mechanism by which sonopermeation can enhance cancer treatment.

## 2 Materials and methods I

### 2.1 Description of the mathematical model

A schematic representation illustrating the various components of the tumor, which have been incorporated into the mathematical model alongside their interrelations, is presented in **Fig 1**, while a presentation of the model’s equations, assumptions, and foundational principles can be found in the Supporting Information (SI) Appendix.

**Fig 1.**
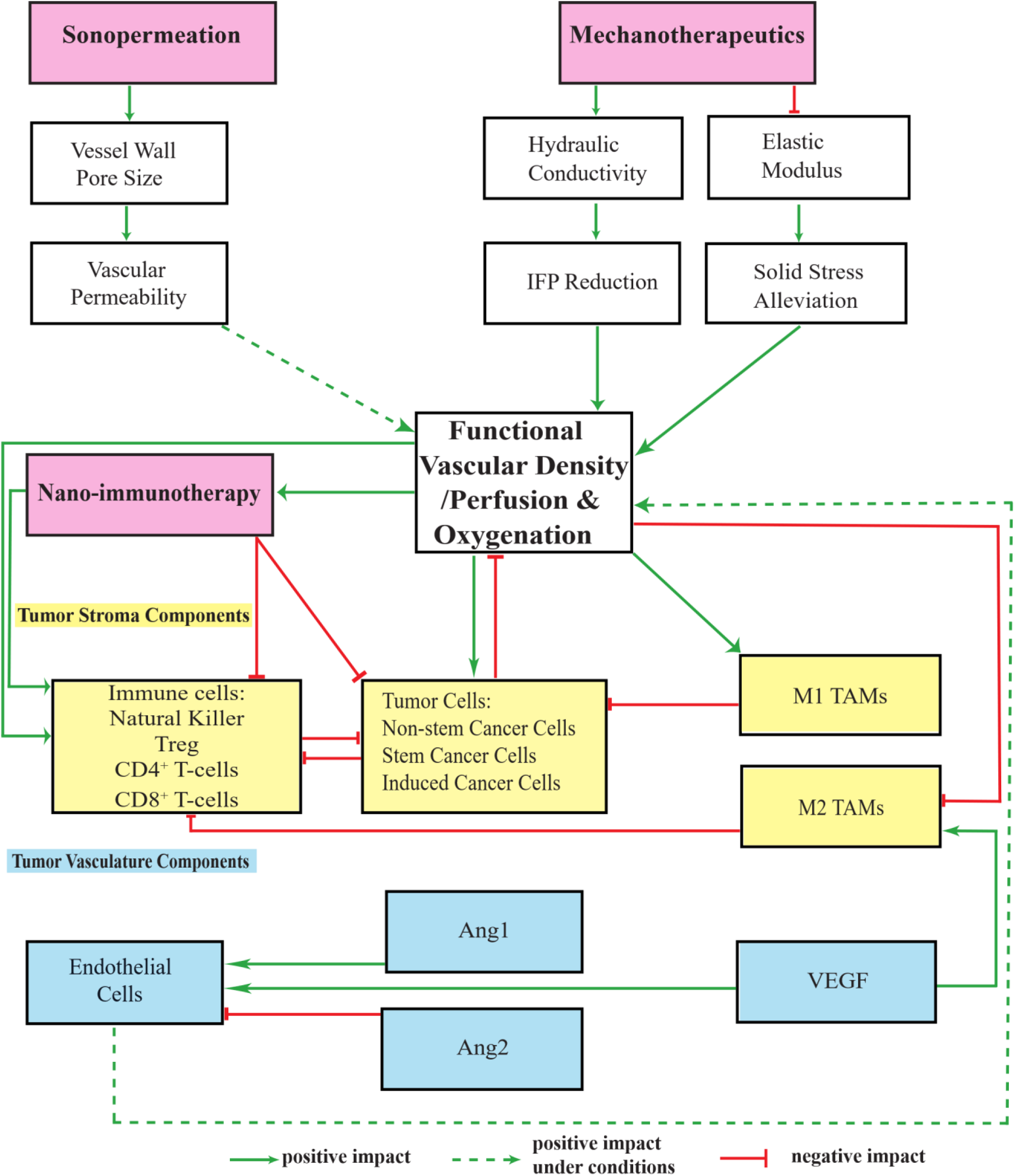
Schematic representation of the components of the mathematical model and their interrelations. The model incorporates diverse populations of cells (illustrated as yellow boxes), tumor angiogenic factors (illustrated as blue boxes), and various therapeutic modalities, each exerting distinct influences within the Tumor Microenvironment (illustrated as pink boxes). This detailed illustration explains the interactions amongst each individual model component, as well as the various potential combinations of these model components, thereby revealing the effects that these interactions have on functional vascular density/perfusion, and oxygenation levels within the examined system. These effects may be classified as positive, negative, or even positive under conditions.

The model accounts for the intricate interactions among various components (**Fig 1**) that are known to play pivotal role in the multifaceted process of tumor progression, such as diverse populations of i) tumor cells: non-stem cancer cells (CCs), stem cancer cells (SCCs), and treatment-induced cancer cells (ICCs) [41], ii) immune cells: NK cells, CD8^+^ T-cells, CD4^+^ T-cells, regulatory T-cells (Treg) and Tumor Associated Macrophages (TAMs), iii) components of the tumor vasculature: Endothelial Cells (ECs), Angiopoietins (Ang), and the Vascular Endothelial Growth Factor (VEGF). The model further accounts for the degree of tumor perfusion, oxygenation and drug delivery (i.e., nano-immunotherapy).

Sonopermeation facilitates an increase in the vessel wall pore size, thereby enhancing vascular permeability, which might augment functional vascular density [28]. The influence of mechanotherapeutics is essential in impacting both the fluid phase and the solid phase of the TME. With respect to the fluid phase, the application of mechanotherapeutics results in a significant elevation of the tumor hydraulic conductivity, which subsequently leads to a decrease in the Interstitial Fluid Pressure (IFP); this decrease, in turn, contributes to an increase in perfusion [6, 24]. As far as the solid phase is concerned, mechanotherapeutics play a critical role in inducing a reduction in the elastic modulus, which ultimately results in the alleviation of solid stress; this alleviation of stress promotes vessel decompression, leading to an increase in functional vascular density/perfusion [6, 24, 25].

An improvement in functional vascular density enhances the efficacy of nano-immunotherapy, which consequently leads to a more effective suppression of non-stem cancer cells, stem-like cancer cells and induced cancer cells [6]. The enhanced concentration levels associated with nanotherapy increase the immunogenic cell death [43, 44] and lead to an improvement in the efficacy of immunotherapy [15, 39],which in turn results in an elevated ratio of CD4^+^/ CD8^+^ T-cells [45]. Through the synergistic effects of nano-immunotherapy, there is a promotion in the recruitment of effector CD8^+^ T-cells, while simultaneously there is a marked reduction in the frequency of regulatory T cells (Tregs), thus fostering a more effective anti-tumor immune response [46]. Furthermore, an increase in the oxygenation not only reinforces the populations of tumor and immune cells but also facilitates the polarization of tumor-associated macrophages (TAMs) from an immune-suppressive M2-phenotype to an immune-activating M1-phenotype [40, 47, 48]. The proliferation of tumor cells leads to significant compression of the surrounding vasculature, which in turn results in a decrease in perfusion and a consequent inactivation of immune cells within the TME [9, 49]. The enhancement of immune cell proliferation boosts the efficacy of tumor cell eradication. M1-like TAMs exert a substantial tumoricidal effect on tumor cells, whereas the M2-like TAMs inhibit the activity of immune effector cells, resulting in the inactivation of immune responses and promoting an immunosuppressive environment [50, 51].

Regarding the tumor vascular components: The process of angiogenesis, which is a fundamental mechanism crucial for the formation of new blood vessels, is triggered through the proliferation of endothelial cells that form the vessels, thereby augmenting the overall perfusion [41]. The elevated levels of VEGF is correlated with an increased number of M2-like TAMs, as well as an elevated proliferation rate of ECs [52]. Furthermore, the presence of high concentrations of Ang2 destabilize existing vessels by diminishing the production of ECs, a phenomenon that is inhibited by Ang1, which contributes in stabilizing vessels and promoting the production of endothelial cells [53, 54].

### 2.2 Transport of Drugs

#### 2.2.1 Transport of Nanotherapy

We postulated that the delivery of nanotherapy exists in three separate states: the nanoparticle carrier containing the chemotherapy (c_n_), the chemotherapeutic agent free to travel in the interstitial space (c_f_) and the chemotherapeutic substance internalized by cells (c_int_) [55]. Therefore, the transport of the drug within the interstitial space can be represented as [56]:

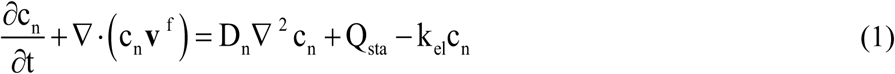

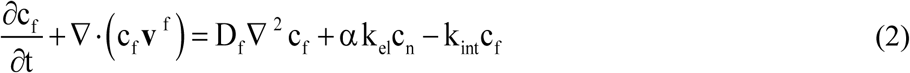

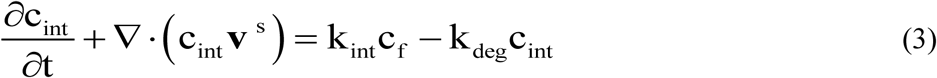

where D_n_ and D_f_ denote the diffusion coefficients of the nanoparticle and chemotherapy within the tumor interstitial space, respectively. The variables k_el_, k_int_ and k_deg_ represent the rate constants for the chemotherapy release, internalization of the drug by the cells, and the degradation rate of the chemotherapeutic agent. Furthermore, α is the number of chemotherapy molecules contained in the nanocarrier and v^f^ and v^s^ are the velocities of the fluid and solid phase, respectively. More information regarding v^f^ and v^s^ is detailed in the SI Appendix, Equations (S18-S26). In the present study, the specific type of nanotherapy employed is Doxil. The term Q_sta_ on the right side of Eq. (1) denotes the transport of the nanocarrier across the tumor vessel wall and it is defined by Starling’s approximation as [56]:

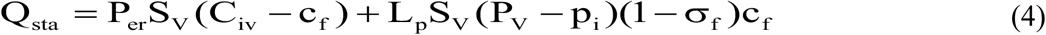

where L_p_ denotes the hydraulic conductivity of the vessel wall, C_iv_=exp(-(t-t_0_)/k_d_) represents the vascular concentration of the administered drug, which is indicative of a bolus injection, with t_0_ being the time of drug administration and k_d_ denoting the blood circulation decay, while σ_f_ is the reflection coefficient. The vascular conductivity L_p_ is determined as a function of the vessel wall pore radius and the parameters P_er_ and σ_f_ are considered as a function of the ratio of the radius of the drug to the radius of the pores of the vessel wall [57]. A detailed description of the methodology and calculations employed in deriving these values can be found within the SI Appendix.

#### 2.2.2 Transport of Immune Checkpoint Antibodies

Immunotherapy is integrated into our mathematical framework through the incorporation of immune checkpoint antibodies, which, in certain therapeutic modalities, can be concurrently utilized to enhance treatment efficacy [41]. In the model, the effect of anti-PD-1 immune checkpoint inhibition is conceptualized as an augmentation in the source term of CD8^+^ T-cells, i.e., the term σ_Τ8_ [41].

Furthermore, the anti-PD-1 antibody is also integrated in our computational model as a free pharmacological agent c_f_i__, and this incorporation is illustrated through the mathematical representation provided in Equation (5).

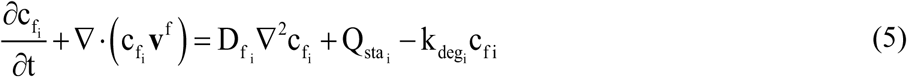

where D_f_i__ denotes the diffusion coefficient of the immune checkpoint antibody within the tumor interstitial space, k_deg_i__ represents the rate constant for the degradation of the anti-PD-1 antibody and v^f^ is the interstitial fluid velocity. More information regarding v^f^ is given by SI Appendix, Equation (S23). The term Q_sta_i__ on the right side of Eq. (5) denotes the transport of the immune checkpoint antibody across the tumor vessel wall and it is defined by Starling’s approximation. It is crucial to recognize that Equation (4), is similarly applied in the context of the transport dynamics concerning the anti-PD-1 antibody, which is capable of moving through the interstitial space.

## 3 Materials and methods II

### 3.1 Modeling the Effects of Mechanotherapeutic Ketotifen

Within the framework of this mathematical model, the incorporation of the mechanotherapeutic ketotifen has been integrated to facilitate the alleviation of stress within solid tumors, thereby leading to a consequential enhancement in the overall efficacy of nano-immunotherapy [6]. The effects associated with ketotifen are a reduction in tumor stiffness, coupled with a notable increase in vascular perfusion, thus enhancing the overall functionality of blood vessels. Experimental data indicate that, three days upon the administration of ketotifen, there is a significant reduction in tumor stiffness by approximately 50% [6, 24, 25]. Furthermore, ketotifen effectively reduces interstitial fluid pressure, facilitate improved tumor perfusion and markedly augmenting the efficacy of drug delivery [16]. In our model, we simulate the effects of ketotifen by reducing the tumor’s shear modulus and increasing the hydraulic conductivity, which in turn leads to a significant reduction in interstitial fluid pressure and increase in functional vascular density.

More specifically, upon the administration of ketotifen, there is a linear decrease in both the shear modulus and the bulk modulus within the tumor tissue in half, which can be quantitatively assessed in relation to the baseline values of these mechanical parameters prior to treatment [24]. Concurrently, the hydraulic conductivity exhibits a linear increase of two orders of magnitude when compared with the initial value of this parameter in the absence of ketotifen intervention. The precise numerical values that are relevant to hydraulic conductivity, k_th_, shear modulus, μ, and bulk modulus, k, concerning the host tissue, the tumor tissue and the tumor tissue with the effect of ketotifen are illustrated in Table S1, SI Appendix.

### 3.2 Incorporation of Sonopermeation

Regarding the phenomenon of sonopermeation, the effect of acoustic pressure on vessel wall pores size has been added in the mathematical model. The process of sonopermeation is known to induce an increase in the size of the pores within the vessel wall, thereby augmenting vascular permeability, which in turn leads to a significant improvement in the functional density of the vascular network. Specifically, it has been demonstrated that sonopermeation possesses the capability to enlarge cell pores, with average dimensions that can vary significantly from 100 nanometers to 1.25 micrometers, and it is noteworthy that the generation of larger sonopermeation pores is positively correlated with an increase in acoustic pressure or an extension of treatment duration of the sonopermeation [58]. By correlating the empirical observations obtained from the aforementioned experimental study with the parameters delineated within our mathematical model, particularly in relation to the influence of acoustic pressure on the size of pores within the vessel wall, we have developed the following second-order polynomial function:

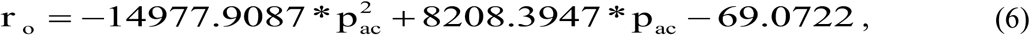

The mathematical expression outlined in Equation (6) describes the effect of acoustic pressure, denoted as p_ac_, on the radius of the pore size within the vessel wall, represented by the variable r_o_, when sonopermeation is applied.

The integration of the acoustic pressure into our mathematical model is accomplished via the subsequent relationship:

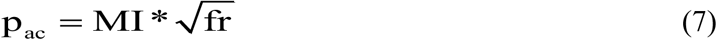

The acoustic pressure, p_ac_, is quantitatively characterized by the mathematical relationship in which the Mechanical Index of the transducer, MI, is multiplied by the square root of the frequency used for sonopermeation, fr, of the wave that has been transmitted [59, 60].

### 3.3 Solution methodology

To effectively model tumor growth, it is presumed that the tumor has a spherical configuration, surrounded by a normal tissue of cubic shape. The cubic host domain, which serves as the spatial environment for the tumor growth, is designed to be two orders of magnitude larger, thereby reducing any potential boundary effects that could interfere with the progression of the tumor. Due to symmetry present in the system under investigation, it is deemed sufficient to consider only one eighth of the entire system for analysis. The clarification of the boundary conditions that have been incorporated in this study is depicted in SI Appendix, Fig S1. In particular, the boundary conditions relevant to the conservation of both the stress and displacement fields, along with the concentration levels of oxygen as well as the immune-nanotherapeutic agents at the interface between the tumor tissue and the adjacent healthy tissue, are applied automatically by the software.

The system of equations that comprise the mathematical model was solved using the commercial finite elements software COMSOL Multiphysics (COMSOL, Inc., Burlington, MA, USA), using the Solid Mechanics, Transport of Diluted Species, Convection-Diffusion Equation and Domain ODEs and DAEs Physics. The computational domain is composed of 6015 finite elements and 51628 degrees of freedom; furthermore, the time-dependent solver that is employed to derive the solutions to the equations governing this model is the PARDISO algorithm.

## 4. Results

### 4.1 Comparative Analysis of Mathematical Model Predictions with Experimental Data

To evaluate the robustness of our mathematical framework and to justify the parameter values employed within the model, we conducted a validation analysis between model predictions with published experimental data [6]. *In vivo* experiments on murine sarcoma models were conducted to determine optimal conditions for combining mechanotherapeutics and ultrasound sonopermeation, in order to improve perfusion and nano-immunotherapy effectiveness [6]. The findings derived from the experimental study revealed that the incorporation of the anti-histamine ketotifen as a mechanotherapeutic with sonopermeation reduced mechanical forces by lowering collagen and hyaluronan levels by 50%, thus reshaping the tumor microenvironment. The combined effects of ketotifen and sonopermeation not only increased tumor perfusion six times but also improved drug delivery. Consequently, the antitumor effectiveness of the Doxil nanomedicine as well as anti-PD-1 immunotherapy was significantly enhanced.

The therapeutic regimen implemented through mathematical modeling was analogous to the experimental protocol utilized for MCA205 fibrosarcoma and K7M2 osteosarcoma tumors (**Figs 2(A) and 3(A)**). For the experimental protocol regarding MCA205 cancer cells, the daily administration of ketotifen started once the tumors reached an average volume of 100 mm^3^. Following a three-day period of ketotifen administration by which time tumors had reached an average volume of 200 mm^3^. Sonopermeation was then employed to synergistically augment the antitumor efficacy of Doxil that has a size of 100 nm in diameter and the immune checkpoint antibody that was taken to have a size of 12 nm. The combined treatment of sonopermeation and nano-immunotherapy was subsequently repeated after four days. The elastic modulus and perfused area were measured during the experimental procedure on specific days utilizing Shear Wave Elastography (SWE) and Contrast Enhanced Ultrasound (CEUS), respectively.

**Fig 2.**
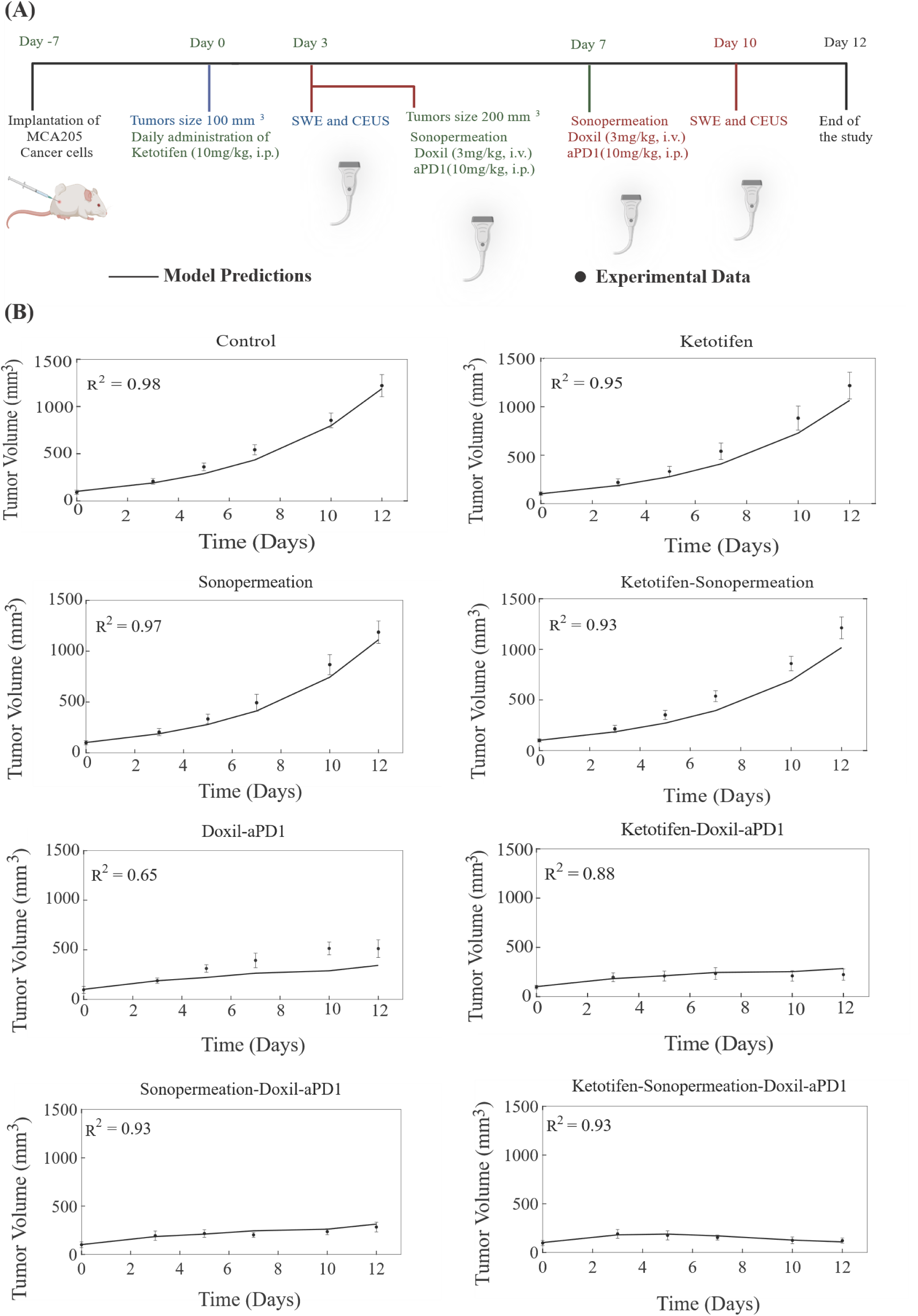
Comparison of model predictions with experimental data of tumor growth for MCA205 fibrosarcoma tumors. **(A)** Experimental treatment protocol followed for MCA205 fibrosarcoma tumors and simulated by the model. Created in <u>BioRender.com</u>. **(B)** Tumor volume growth rates of murine fibrosarcoma cancer cells (dots) and mathematical model predictions (solid lines) for each treatment group. For each case - control, ketotifen, sonopermeation, ketotifen-sonopermeation, Doxil-aPD1, ketotifen-Doxil-aPD1, sonopermeation-Doxil-aPD1 and ketotifen-sonopermeation-Doxil-aPD1- the R-Squared (R²) value has been calculated and depicts the accuracy of mathematical model validations for tumor growth in comparison with experimental findings. aPD1 denotes for anti-PD1 antibody.

**Fig 3.**
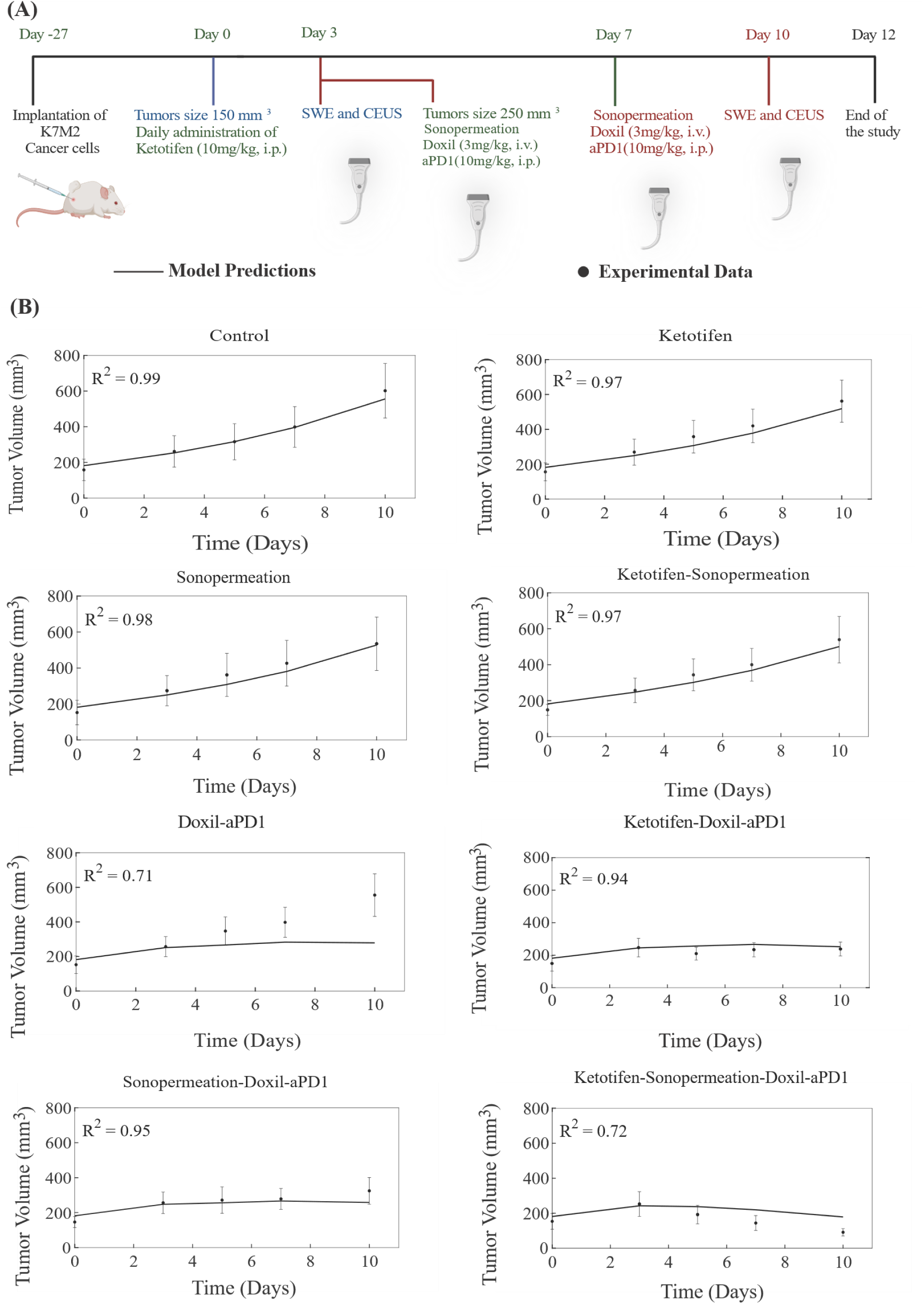
Comparison of model predictions with experimental data of tumor growth for K7M2 osteosarcoma tumors. **(A)** The experimental treatment protocol implemented for K7M2 osteosarcoma tumors and simulated by the model. Created with <u>BioRender.com.</u> **(B)** The tumor volume growth of murine osteosarcoma cells (dots) along with the predictions derived from mathematical modeling (solid lines) for each treatment group. For each case - control, ketotifen, sonopermeation, ketotifen-sonopermeation, Doxil-aPD1, ketotifen-Doxil-aPD1, sonopermeation-Doxil-aPD1 and ketotifen-sonopermeation-Doxil-aPD1-the R-Squared (R²) value has been calculated and depicts the accuracy of mathematical model validations for tumor growth in comparison with experimental findings. aPD1 denotes for anti-PD1 antibody. To elucidate the comparative analysis between the model and experimental data, we further compared model predictions with measurements of perfused area measured with contrast enhanced ultrasound, elastic modulus measured with shear wave elastography and drug concentration measured with fluorescence imaging and IFP measurements (Fig 4). To correlate the dimensionless parameters of the model and the actual measurements obtained from the experiments, the values corresponding to the various measured parameters are presented in relation to the values of the control group, specifically represented as a fold change. It is noteworthy that the predictions derived from the model show a substantial level of agreement with the experimental data concerning the perfused area, the elastic modulus, the levels of drug concentration and the IFP as illustrated in Fig 4. In situations where ketotifen is administered in conjunction with sonopermeation and concurrent administration of ketotifen-sonopermeation-Doxil-aPD1, it is observed that there is a significant enhancement in the perfused area, which in turn leads to an improvement in the functional vascular density Fig 4**(A)** and **4(D)**. Tumors that have undergone a pretreatment regimen involving the administration of ketotifen, exhibit a marked reduction in the tumor elastic modulus alongside an increase in tumor perfusion **Figs 4(B)** and **4(E)**. Moreover, by comparing the concentration levels of the drug with the various experimental groups that have been administered Doxil and aPD1, the predictions generated by our model exhibit a good level of agreement with the experimental data, illustrating that the synergistic effects resulting from the combination of ketotifen and sonopermeation indeed contribute to an enhancement in the drug delivery process (Fig 4**(C)**). Last but not least, the combination of both ketotifen and sonopermeation prior to the injection of nano-immunotherapy results in a substantial decrease in interstitial fluid pressure (IFP) when compared to the group that received only ketotifen or the group that received both ketotifen and nano-immunotherapy (Fig 4**(F)**).

**Fig 4.**
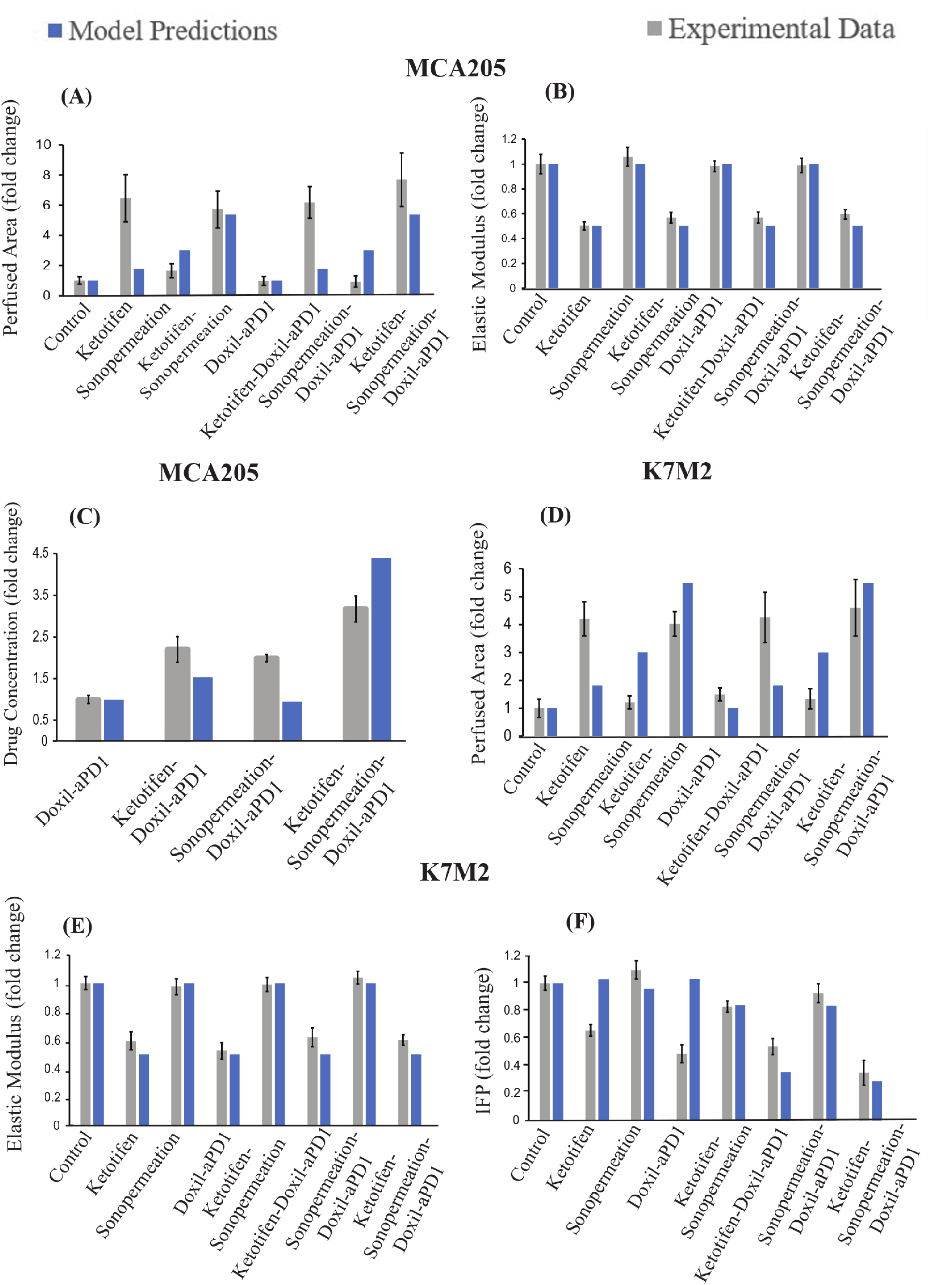
Comparison of model predictions in conjunction with experimental data [6] for a specific time point. The horizontal axis delineates the various treatment groups that were included in the experimental investigations: control, ketotifen, sonopermeation, ketotifen-sonopermeation, Doxil-aPD1, ketotifen-Doxil-aPD1, sonopermeation-Doxil-aPD1, ketotifen-sonopermeation-Doxil-aPD1. The vertical axis (y) for each instance varies between **(A)** Perfused Area, **(B)** Elastic Modulus and **(C)** Drug Concentration for MCA205 fibrosarcoma, **(D)** Perfused Area, **(E)** Elastic Modulus and **(F)** Interstitial Fluid Pressure (IFP) for K7M2 osteosarcoma.

In order to carry out a comparison between the predictions generated by the model and the experimental data, all model parameters were set to baseline values derived independently from relevant studies (SI Appendix, Table S1). The only model parameter that underwent modification in order to adequately align the model’s predictions with the experimental findings [6] was the parameter denoted as k_1_, which serves to quantify the relationship between the proliferation of cancer cells and the concentration of oxygen (SI Appendix, Equation S8). The value assigned to k_1_ was determined such that the predicted final tumor volume produced by the model was the same as the experimental measurement obtained from the control/untreated group, and this value was kept the same across all comparisons of model predictions against the complete set of experimental data derived from all groups participating in the same study. Consequently, it is noteworthy that despite the presence of a considerable number of parameters incorporated within the model—an aspect that is attributed to the mechanistic nature of the model—only a single parameter was subject to variation for each sarcoma type (SI Appendix, Table S2).

The tumor volume estimations derived from mathematical modeling exhibit strong agreement with the experimental data, as monotherapies—namely control solution, ketotifen, sonopermeation, and the combination of ketotifen with sonopermeation—demonstrated no antitumor effects in terms of tumor volume reduction relative to the control group. Conversely, a notable reduction in overall tumor volume was observed for the combinatorial treatment. Our model substantiates that the combination of ketotifen and sonopermeation with nano-immunotherapy significantly improved therapeutic outcomes and particularly suppressed tumor growth (**Figs 2(B) and 3(B)**). A more pronounced delay in tumor growth for both MCA205 and K7M2 sarcomas was recorded in the treatment modality where ketotifen, sonopermeation, Doxil, and anti-PD1 antibody were combined, resulting in effective tumor volume reduction. The predictive capacity of the mathematical model, which aligns closely with experimental observations, can be evaluated using the R-Squared (R²) statistic, which measures the precision of the correlation between experimental data and model forecasts, ranging from 0 to 1, with enhanced precision indicated by R² values approaching 1.

### 4.2 Parametric Analysis

#### Parametric Analysis of Essential Model Variables

The goal of the parametric analysis is to elucidate the specific values of distinct parameters that exert the most significant influence on the phenomenon of sonopermeation and consequently, the resultant therapeutic outcomes that can be achieved through this technique.

Upon the application of sonopermeation, there is a marked enhancement in the pore size, or permeability, of the vessels walls associated with tumor tissues, leading to greater drug infiltration. In order to facilitate a controlled and effective delivery of the therapeutic agent without incurring any detrimental damage to surrounding tissues, it is imperative to meticulously regulate the parameters associated with ultrasound application. In the context of sonopermeation, the ultrasound is typically applied in a pulsed manner to reduce tissue damage from excessive heating and enable microbubble inflow during pulse intervals, especially when bubble destruction is likely. The sinusoidal ultrasound wave is characterized by parameters such as velocity, wavelength, frequency, pressure amplitude, pulse length (burst duration), pulse repetition frequency (PRF), exposure time (duty cycle), treatment duration, and post-sonopermeation effects [27, 61–63]. The parameters commonly utilized for sonopermeation and subsequent drug delivery exhibit considerable variability across different research studies, encompassing frequency ranges from 0.5 to 3 MHz, pressure levels fluctuating between 0.05 to 2 MPa, and total treatment durations that can span from mere seconds to several hours [28, 62, 64–70]. Furthermore, the fundamental effect of sonopermeation persists for a duration ranging from 4 to 24 hours, and after the lapse of 24 hours, it reveals no substantial effect [71, 72].

In view of the previously mentioned parameters, we opted to integrate and simulate within our mathematical model the frequency employed for sonopermeation, the acoustic pressure that we correlate with the mechanical index (MI) through the Equation (7) and the effect of treatment related to sonopermeation. To perform parametric analysis, we varied values of frequency and MI within the ratio given previously and check treatment efficacy for different time points that sonopermeation effect occurs.

**Fig 5** depicts the impact exerted by the frequency of sonopermeation on various critical parameters, including the pore radius, interstitial fluid pressure, tumor volume, functional vascular density, and the concentration of the administered drug. A detailed examination reveals that when the sonopermeation frequency is situated within a specific range, particularly around the value of 0.5 MHz, the resultant data indicate that these conditions yield optimal outcomes across all the aforementioned parameters that are represented within this figure.

**Fig 5.**
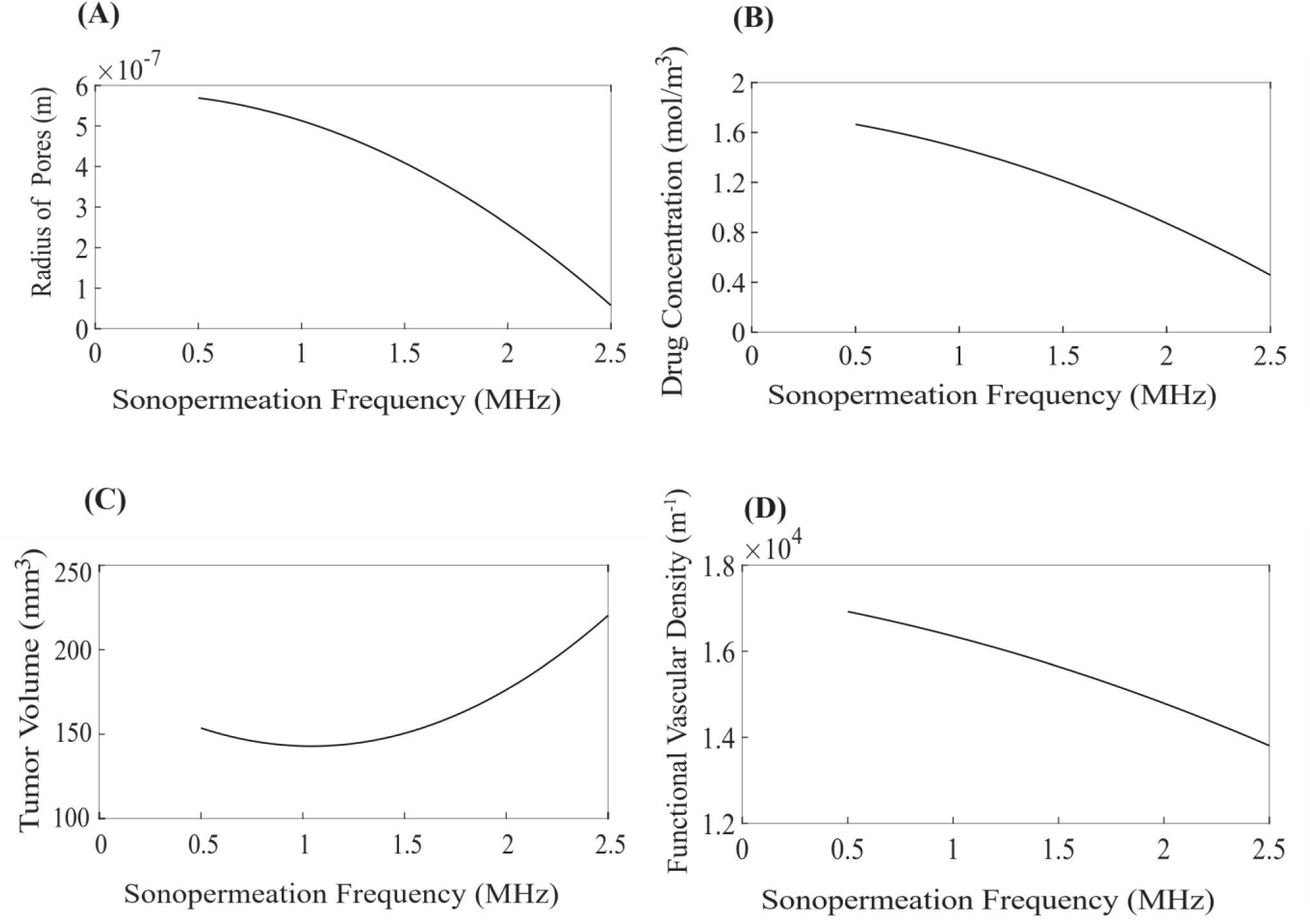
Impact of sonopermeation frequency on the following parameters: **(A)** pore radius (m), **(B)** drug concentration (mol/m^3^), **(C)** tumor volume (mm^3^) and **(D)** functional vascular density (m^−1^). The values corresponding to the various model parameters were computed at a position that is equidistantly located between the central region of the tumor and its peripheral boundaries.

In **Fig 6**, we examined the impact of the mechanical index of the transducer on the pore radius, interstitial fluid pressure, tumor volume, functional vascular density, and the concentration of the relevant drug. An evaluation of the data suggests that the most favorable results become apparent when the mechanical index is approximately 0.17, as this value correlates positively with all the parameters that are illustrated.

**Fig 6.**
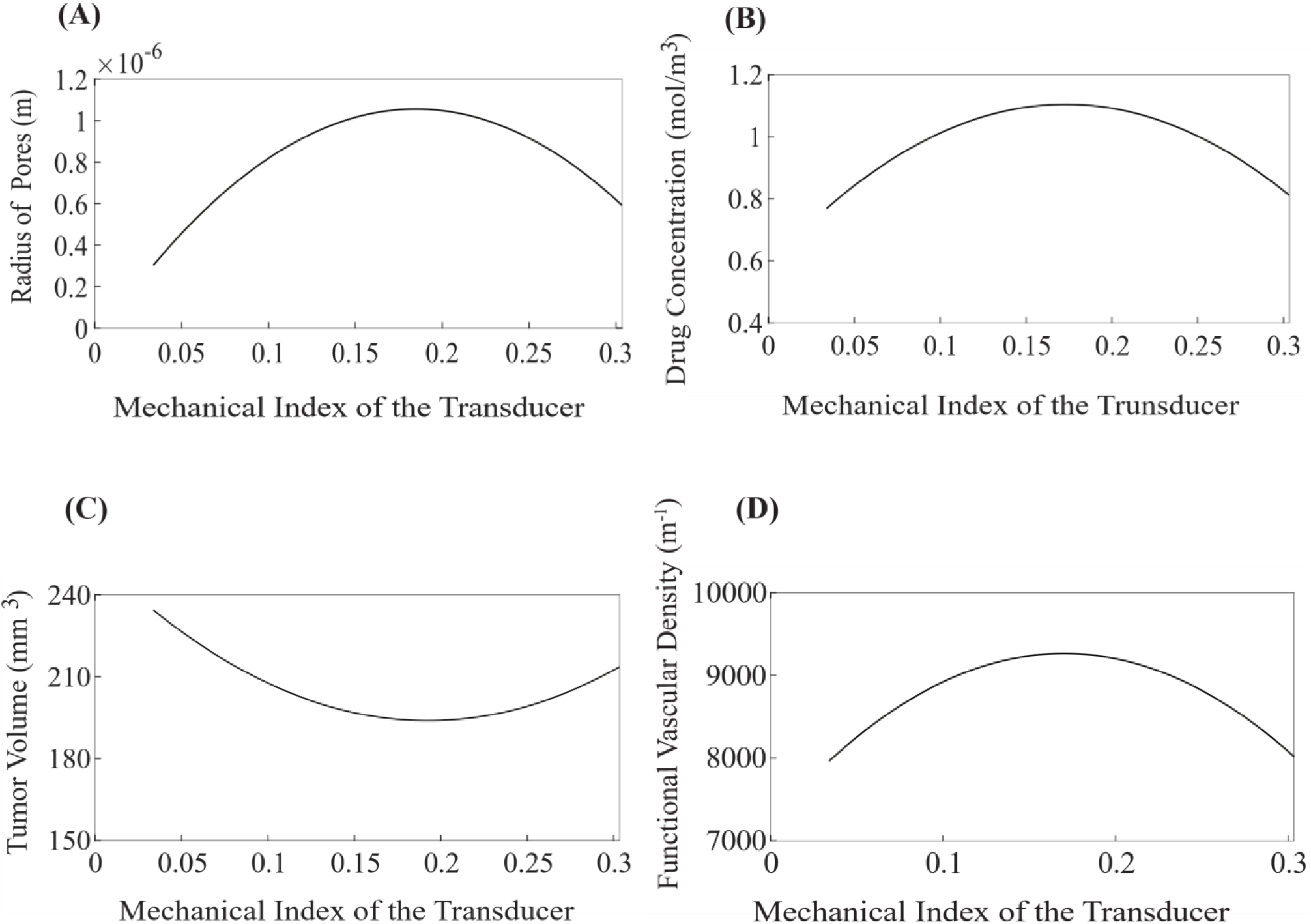
Impact of mechanical index of the transducer on the following parameters: **(A)** pore radius (m), **(B)** drug concentration (mol/m^3^), **(C)** tumor volume (mm^3^) and **(D)** functional vascular density (m^−1^). The values corresponding to the various model parameters were computed at a position that is equidistantly located between the central region of the tumor and its peripheral boundaries.

In conclusion, it can be stated that when the frequency of sonopermeation (fr) approaches 0.5 MHz and the mechanical index (MI) fluctuates around the value of 0.17, the corresponding acoustic pressure (p_ac_) is found to be 0.24 MPa and 0.25 MPa, respectively.

As illustrated in **Fig 7**, it becomes clear that the optimal combination for minimal tumor volume and maximum drug concentration, is attained when the frequency is defined in the vicinity of 2.5 MHz, the mechanical index (MI) is established in the vicinity of 0.17, and consequently the acoustic pressure is calculated as 0.27 MPa. Thus, it is evident that the most favorable outcomes manifest when the acoustic pressure is maintained within the critical range of 0.24 to 0.27 MPa, as this condition appears to optimize the critical parameters being analyzed. The maintenance of acoustic pressure within the specific and critical parameters ranging from 0.24 to 0.27 MPa yields the most favorable and effective outcomes, as this particular condition seems to enhance the concentration of the administered pharmaceutical agents and thus, antitumor effects. Furthermore, it is worth noting that it does not appear to be a difference between the durations of the effect of sonopermeation set at 24 hours compared to those in 72 hours (**Figs 7(E-H)**), which serves to reaffirm the experimental observation of the effect of sonopermeation predominantly lasts for a maximum of 24 hours. As evidenced by the data presented in **Figs 7(E) and 7(F)**, one can observe a remarkable reduction in tumor growth that occurs alongside the elevated concentration of the drug, in addition to the notable increased in vascular density and the optimization of pore size. In **Fig 7(H)** the drug concentration is higher than in the **Fig 7(F)** but the tumor volume is approximately the same (**Figs 7(E)** and **7(G))**.

**Fig 7.**
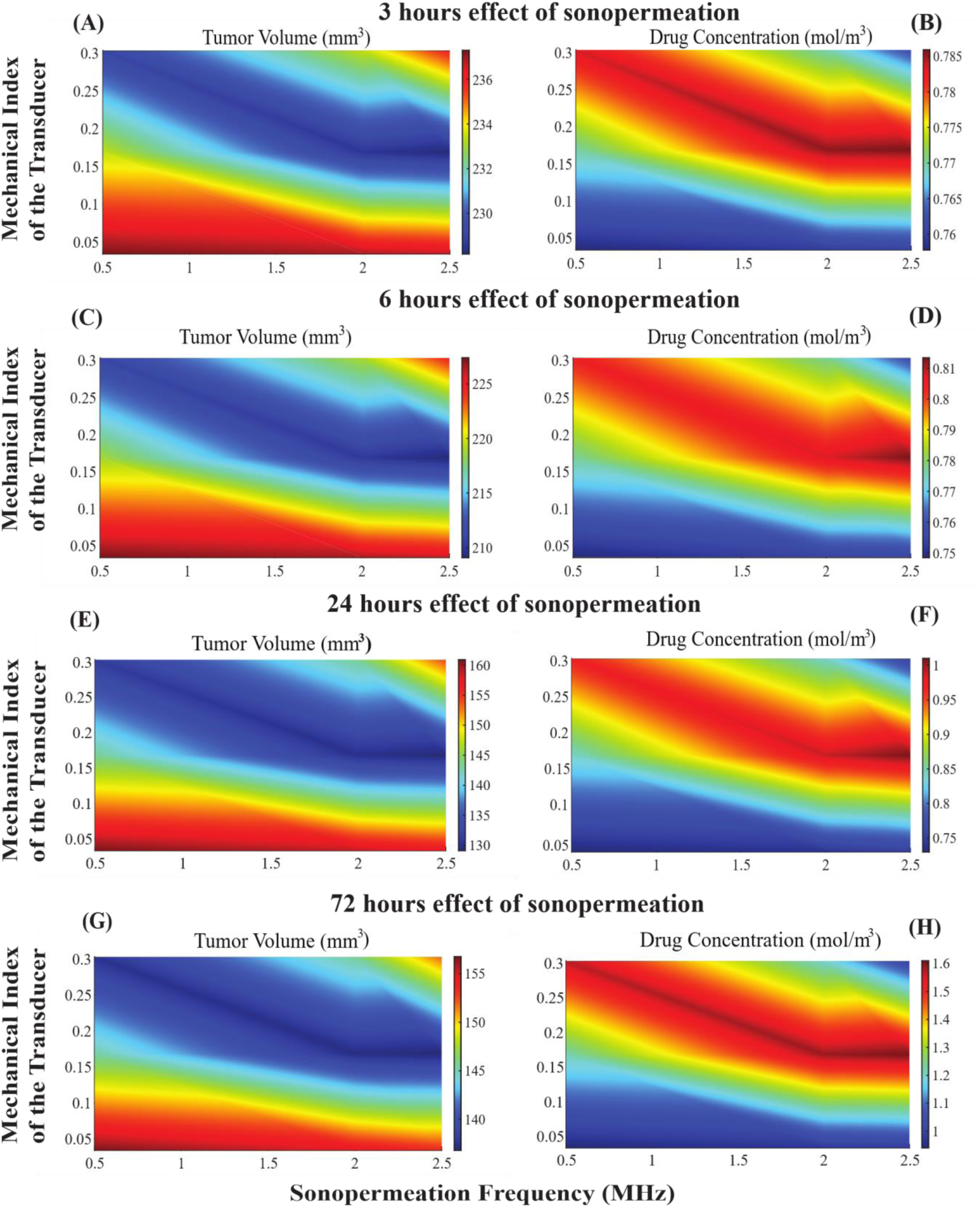
The influence of varying sonopermeation frequencies, in conjunction with diverse values of the mechanical index (MI) of the transducer utilized for the application of sonopermeation, on the synergistic effects of combined therapy is analyzed. Phase diagrams depict the impact of treatments on tumor volume **(A)**, **(C)**, **(E)**, and **(G)** as well as drug concentration **(B)**, **(D)**, **(F)**, and **(H)** across different time frames of the effect of sonopermeation. Specifically, **(A)** and **(B)** correspond to a 3-hours **(C)** and **(D)** to a 6-hours, **(E)** and **(F)** to a 24-hours effect of sonopermeation and **(G)** and **(H)** to a 72-hours effect of sonopermeation.

## 5. Discussion

In this study, we have formulated a mechanistic mathematical framework that integrates the synergistic application of mechanotherapeutics and sonopermeation, with the primary objective of realizing the highest possible efficacy of nano-immunotherapy in the treatment of cancer. The model we developed encompasses the diverse interactions that occur among various categories of cancer cells, immune cells, tumor associated macrophages, endothelial cells, tumor angiogenic factors and an array of different therapeutic modalities, each of which exerts its unique and distinct influences within the complex domain of the tumor microenvironment. Our model represents a significant advancement over prior research endeavors, as it incorporates the effects of sonopermeation alongside the mechanotherapeutic agent ketotifen, which is known for its therapeutic potential. The complexity of our model can be substantiated by the favorable alignment of the predictions generated by the mathematical model with the experimental results obtained from in vivo studies on two distinct sarcoma cell lines that exhibit varying growth rates, specifically one characterized by a rapid growth rate (the fibrosarcoma MCA205) and another that demonstrates a comparatively slower growth rate (the osteosarcoma K7M2). The experimental data we have collected provide compelling evidence that the mechano-modulation of the tumor microenvironment, achieved through the combined application of mechanotherapeutics and sonopermeation, can result in multiplicative synergistic effects that significantly enhance perfusion and improve overall therapeutic outcomes [6]. These observations are validated with a good degree of precision through our mathematical model. A good correlation was observed between the computational values produced by the model and the actual experimental measurements of critical parameters, such as tumor volume, functional vascular density, elastic modulus, interstitial fluid pressure, drug concentration which are essential to reinforce the overall accuracy and reliability of the mathematical framework employed.

The parametric analysis, which examined a variety of parameters, provided optimal guidelines that are essential for the effective implementation of sonopermeation. These parameters include, the mechanical index of the transducer utilized, the specific frequency at which sonopermeation is conducted, the acoustic pressure applied during the procedure, and the duration over which the effect of sonopermeation is exerted. More specifically, the analysis delineates a precise range of values for these critical parameters within which the efficacy of sonopermeation is markedly enhanced, thereby yielding the most favorable and effective results.

This particular model has certain limitations especially in relation to the application of the effect of sonopermeation within the framework of our model. Sonopermeation could improve the overall efficacy of drug delivery by leveraging additional mechanisms that are not explicitly included in the current model framework. For instance, the interaction of acoustic waves and the resultant shear wave stresses, that are exerted upon endothelial cells by the action of microbubbles has been defined in accordance with prior studies [36, 73, 74]. Furthermore, the effects of sonopermeation on the extracellular matrix would also be integrated into the model. It is worth mentioning that the application of shear stress has been shown to reduce apoptosis in endothelial cells, thereby enhancing functional vascular density [75]. The previously mentioned limitations are expected to influence the quantitative aspects of the predictions generated by the model, while the fundamental conclusion derived from this comprehensive study remains unaffected.

## Acknowledgments

This project received funding from the European Research Council (ERC) under the European Union’s Horizon 2020 and Horizon Europe research and innovation programme (grant agreement nos. 863955 and 101069207 to TS and 101076425 to FM).

## Author contributions

All authors contributed to conceptualization, methodology, validation, formal analysis, investigation, resources, data curation, writing – review and editing and visualization of the article. Marina Koutsi was responsible for writing the original draft. Fotios Mpekris and Triantafyllos Stylianopoulos were responsible for supervision and project administration and funding acquisition.

